# Betrayal and unfairness are linked through insula-valuation network dynamics

**DOI:** 10.1101/2025.11.20.689349

**Authors:** Jacob M. Stanley, Derrick A. Dwamena, James B. Wyngaarden, Melanie C. Kos, Johanna Jarcho, Dominic Fareri, David V. Smith

## Abstract

Trust and fairness sustain cooperation, but social neuroscience has largely studied violations of these social norms in isolation. We asked whether neural sensitivity to betrayal carries forward into the evaluation of unfairness in a separate social exchange. In 132 adults who completed fMRI during the Trust Game and Ultimatum Game, betrayal and unfairness both recruited overlapping voxels within left anterior insula. Yet, individual differences in regional activation were uncorrelated across tasks, arguing against a simple shared-response account. Instead, cross-task structure emerged in connectivity: stronger anterior insula responses to social versus nonsocial betrayal predicted weaker anterior insula–ventromedial prefrontal coupling during social versus nonsocial fairness evaluation. These findings suggest that betrayal and unfairness are linked less by identical regional responses than by how salience signals engage valuation circuitry across contexts. By bridging two canonical social exchange tasks within the same individuals, this study reveals a network-level architecture for generalizing sensitivity to interpersonal norm violations.

## Introduction

Fairness and trust are cornerstones of social interaction, shaping cooperation, resource distribution, and relationship maintenance. Both represent deeply valued social norms, and violations of these norms evoke robust behavioral and neural responses. The Ultimatum Game (UG) and the Trust Game (TG) are two influential paradigms for studying these processes, providing a controlled setting to examine how individuals respond to unfairness, betrayal, and reciprocity during social economic exchanges. Although behavior across economic exchange games is correlated (Chapman et al., 2023), whether these shared tendencies reflect common neural organization remains unknown. By measuring UG and TG responses in the same participants, the present study tests whether sensitivity to one form of norm violation carries across contexts at the level of behavior, regional activation, and task-modulated connectivity.

Prior UG research suggests that unfair offers engage both affective and deliberative processes. From a dual-process perspective, norm violations can trigger rapid affective responses while also recruiting evaluative mechanisms that weigh material gain against punishment of unfairness (Evans, 1984; Kahneman, 2011; Sanfey et al., 2003; Fehr & Camerer, 2007). The anterior insula (aIns) is especially relevant to the affective component of this response. Broadly, the aIns is thought to integrate interoceptive information with salience detection, supporting awareness of bodily and affective states that signal when events are personally or socially significant (Craig, 2009; Menon & Uddin, 2010). Consistent with this account, UG studies have repeatedly linked aIns activity to unfair offers, perceived injustice, and inequity aversion (Sanfey et al., 2003; Gabay et al., 2014; Tabibnia, et al., 2008). The aIns may therefore encode the aversive salience of unfair treatment and interact with valuation-related regions such as ventromedial prefrontal cortex (vmPFC; Bartra et al., 2013; Dennison et al., 2022; Hampton et al., 2006; Jenkins et al., 2008; Ruff & Fehr, 2014; Smith & Delgago, 2015), providing a pathway through which socially salient norm violations can shape downstream evaluation and choice.

Although the Trust Game differs structurally from the Ultimatum Game, requiring an initial decision to invest before observing partner behavior, it similarly involves evaluating whether a social partner has violated expectations. This raises the question of whether affective responses to betrayal recruit the same salience-detection mechanisms implicated in responses to unfairness. Prior work has shown that the caudate nucleus tracks partner trustworthiness (Delgado et al., 2005; King-Casas et al., 2005), the amygdala contributes to the development and expression of interpersonal trust (Koscik & Tranel, 2011), and medial prefrontal regions respond more strongly to human than computer partners (McCabe et al., 2001). Critically, trust violations are salient social events, making the anterior insula a plausible candidate for detecting and representing betrayal-related affective significance. Indeed, anterior insula activity has been linked to reciprocity-related behavior in the TG (van den Bos et al., 2009), suggesting that this region may contribute not only to responses to unfairness in the UG, but also to the evaluation of socially meaningful outcomes in trust-based exchange. Together, these findings raise the possibility that the anterior insula contributes a common salience signal across betrayal and unfairness, while leaving open whether that commonality appears in regional activation, connectivity with valuation systems, or both.

Despite these parallels, research has rarely tested UG and TG responses within the same individuals. This separation leaves open whether betrayal and unfairness are linked by common behavioral tendencies, shared regional responses, or network-level interactions between salience and valuation systems. We therefore asked three questions. First, do behavioral responses to unfairness in the UG show sensitivity to fairness and social context? Second, do betrayal in the TG and unfairness in the UG recruit overlapping anterior insula responses, and are those responses correlated across individuals? Third, if regional activation does not provide a direct bridge between tasks, do cross-task relationships emerge in anterior insula connectivity with valuation-related regions during fairness evaluation?

## Results

To answer these questions, we first examined behavior in the UG. Illustrated in Figure 1, the UG tasks participants to accept or reject monetary offers from a partner’s endowment, which provides a lens through which to examine fairness processing by comparing acceptance rates across levels of offer equity. We then tested whether occurrences of UG unfairness and TG betrayal recruited overlapping anterior insula responses. The TG (Figure 1) tasks participants to choose an amount to invest in a partner that may or may not elect to share the return on their investment, allowing a measure of neural responses toward unreciprocated trust, i.e. betrayal, that we hypothesize shares similarities with unfairness processing in the UG. Finally, we asked whether individual differences in betrayal-related anterior insula activity predicted task-modulated connectivity during fairness evaluation.

**Figure 1.**
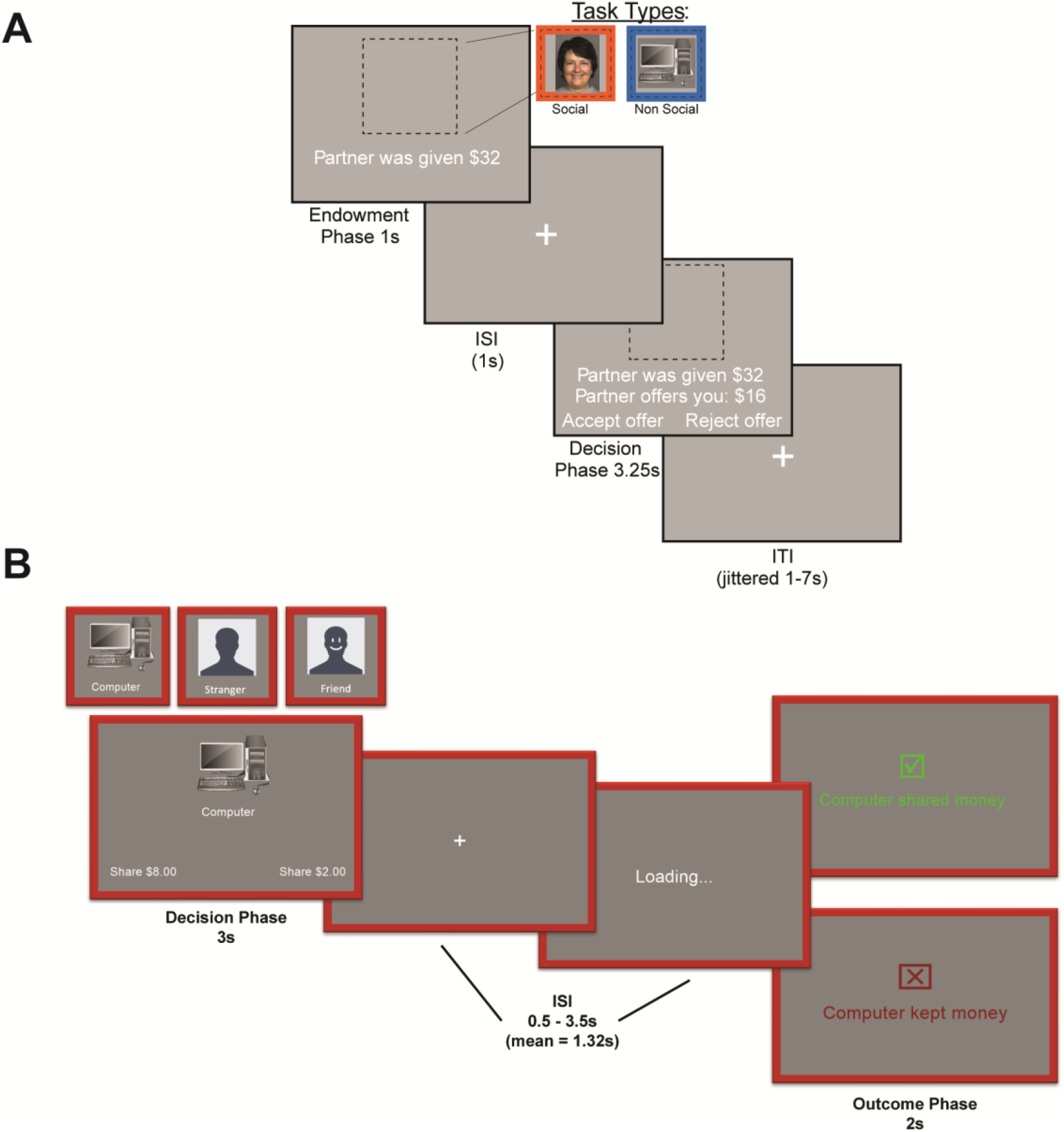
Task Timelines. **A.** Participants in the Ultimatum Game decided to accept or reject offers made by a social or nonsocial partner. If they rejected the offer, neither the participant nor their partner would make any money. Participants were offered between 5% and 50% of the initial endowment. **B**. Participants in the Trust Game play each round with one of three possible partners: a computer, a stranger, or their friend. During the decision phase, participants choose one of two possible amounts to invest in their partner, which is tripled and given to them. The participant keeps the remainder. During the outcome phase, participants see whether their friend chose to share the tripled amount evenly with them or not.

### Acceptance Rates Increase with Offer Fairness

Behavioral responses in the Ultimatum Game were analyzed to quantify acceptance rates as a function of offer fairness and partner type. Trial-level accept/reject decisions were converted to a binary variable (Accept = 1, Reject = 0). For each participant, we computed mean acceptance rates separately for each fairness level (5%, 10%, 25%, 50% of the endowment) and each partner condition (social vs. nonsocial). These subject-level acceptance rates served as the dependent measure in a two-way repeated-measures ANOVA with fairness (4 levels) and partner (2 levels) as within-subject factors.

The analysis revealed a robust main effect of fairness, *F*(3, 393) = 202.85, *p* < .001, η^2^ = .39, indicating that acceptance rates increased strongly as offers became more equitable. In contrast, there was no main effect of partner, *F*(1, 131) = 1.87, *p* = .17, and no interaction of fairness and partner, *F*(3, 393) = 0.14, *p* = .94. Thus, participants responded robustly to fairness of the offer but did not differentiate between social and nonsocial proposers. These results are shown in Figure 2.

**Figure 2.**
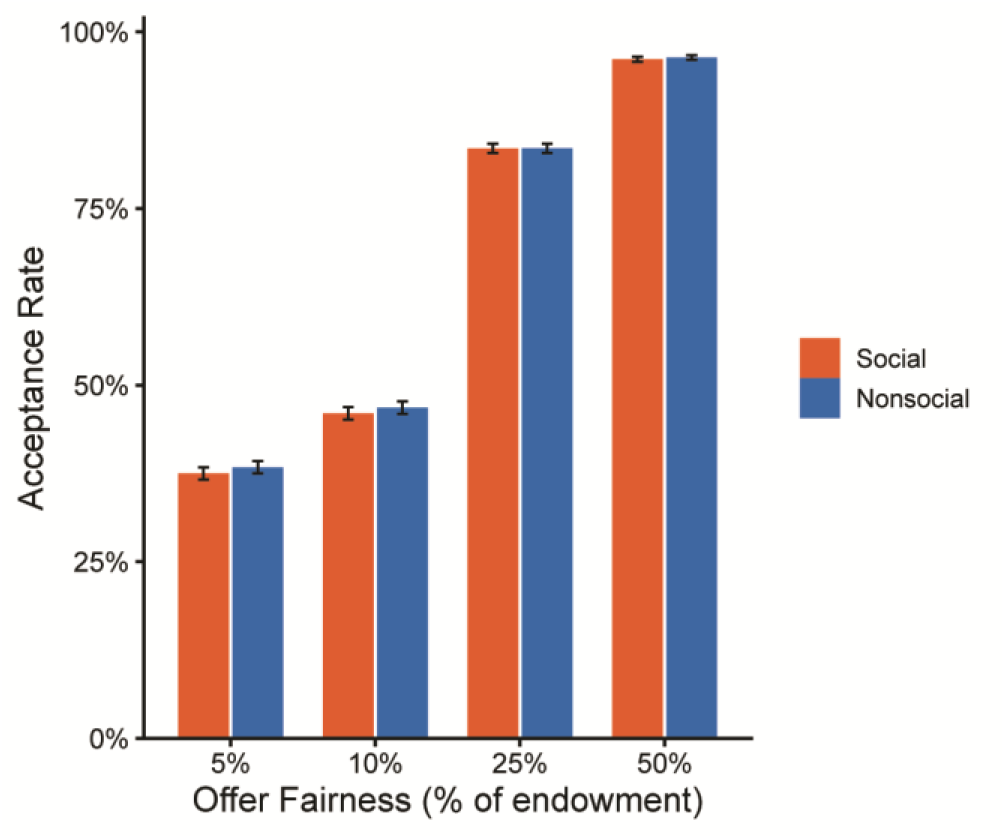
Acceptance Rates as a Function of Offer Fairness and Partner Type. Mean acceptance rates in the Ultimatum Game across four levels of offer fairness for social and nonsocial partners. Error bars represent ±1 standard error of the mean.

### Anterior Insula Activity Tracks Offer Fairness and Trust Betrayal

A whole-brain activation analysis of the Ultimatum Game offer phase revealed that activity in the anterior insula tracked offer fairness. Specifically, the anterior insula showed a significant negative parametric relationship with the percentage of the endowment offered, such that activation decreased as offers became more fair. This effect was observed in the left anterior insula (peak MNI = -34.3, 19.8, -12.6, k = 137 voxels, peak Z = 5.11, cluster corrected *p* < .001). As illustrated in Figure 3, mean signal extracted from this cluster showed lessened responsivity to more equitable offers across both social and nonsocial conditions, consistent with a role for the anterior insula in encoding the relative aversiveness or salience of unfair relative to fair treatment.

**Figure 3.**
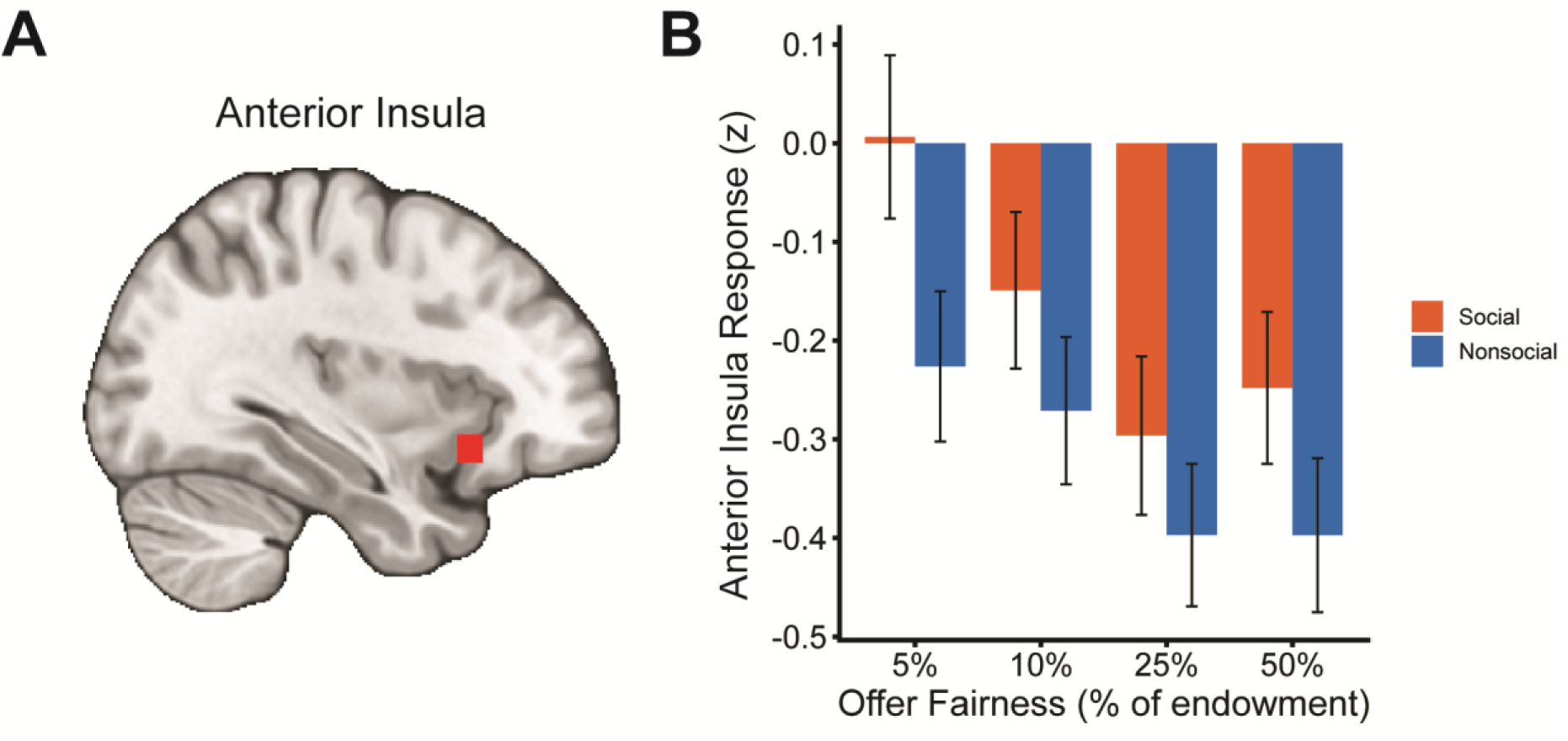
Anterior insula sensitivity to offer fairness in the Ultimatum Game. **A.** Whole-brain parametric modulation analysis of the offer phase revealed a significant cluster in the left anterior insula in which activity decreased as offer fairness increased. **B**. Plot of mean extracted signal from the cluster across offer fairness levels for social and nonsocial conditions, shown for visualization of the effect direction only.

A whole-brain activation analysis of the Trust Game outcome phase revealed that the left anterior insula showed greater activation for trials where trust was violated compared to when trust was reciprocated (peak MNI = -45.1, 22.5, -0.72, k = 746 voxels, peak Z = 7.30, cluster corrected *p* < .001). Mean signal extracted from this cluster showed greater responsivity to outcomes where the partner did not share the multiplied investment compared to outcomes where the partner did share.

Because the anterior insula was found to respond more to unreciprocated trust in the TG and to track offer fairness in the UG, we hypothesized that anterior insula BOLD responses across these two tasks would be correlated within-subjects. To test this hypothesis, we extracted mean beta weights from voxels in the anterior insula that belonged to the intersection of significant clusters in both the defect > reciprocate contrast of the TG and the parametric modulation analysis tracking offer fairness in the UG. However, we did not find a significant association between the BOLD response across the two tasks, *r* = -.05, *p* = .59. This relationship, as well as the overlapping voxels across task, are displayed in Figure 4.

**Figure 4.**
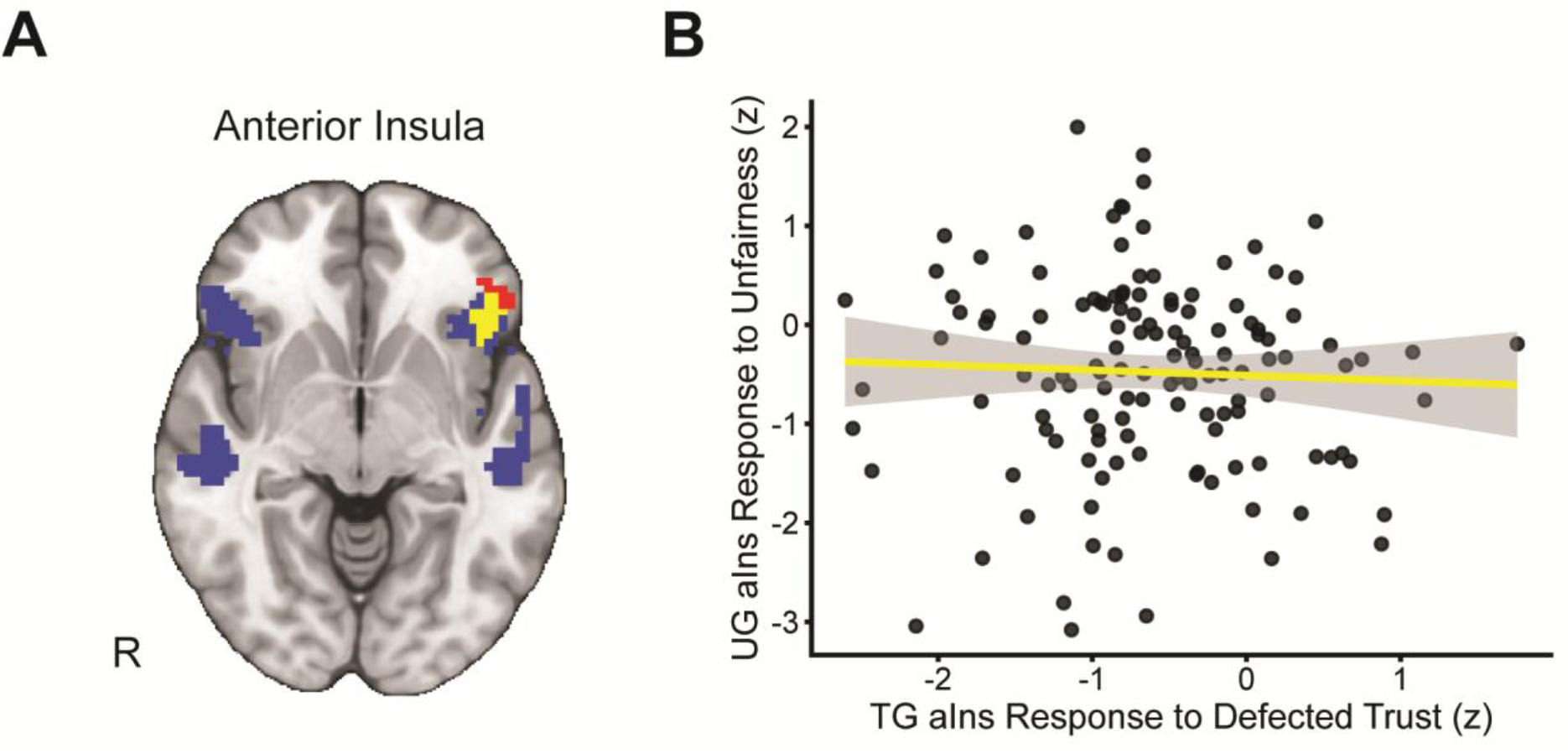
Cross-task anterior insula response to norm violations. **A.** Red voxels are those showing a significant negative correlation with a parametric modulator tracking offer fairness during the UG. Blue voxels are those showing greater activation for trials where trust is defected compared to trials where trust is reciprocated. Yellow voxels are those which are significant for both across task. **B**. Scatterplot showing the relationship between anterior insula response to unfairness in the UG and unreciprocated trust in the TG for the yellow voxels in panel A.

### Trust-related aIns Activity Predicts Fairness-related aIns-vmPFC Connectivity

In order to further examine the cross-task role of the anterior insula, which showed preferential responsivity to defection of trust in the TG and also tracked offer fairness in the UG, we conducted a secondary whole-brain psychophysiological interaction (PPI) analysis for social > nonsocial UG offers using the anterior insula (aIns) as the seed. We included a between-subjects covariate representing each participants’ response defected > reciprocated trust during social > nonsocial offers in the TG within the same aIns seed region. The aIns seed was defined as the binary intersection of voxels that tracked offer fairness in the UG and voxels that selectively responded to betrayal of trust in the TG, represented by the yellow voxels in Figure 4. We identified a significant cluster in the ventromedial prefrontal cortex (vmPFC; peak MNI = -1.9, 41.4, -9.6, k = 18, peak Z = 4.35, cluster-corrected *p* = .024). Connectivity between the aIns and vmPFC for the Social > Nonsocial contrast in the Ultimatum Game was negatively associated with aIns activation for the Social > Nonsocial betrayal contrast in the Trust Game (Figure 5). Thus, individuals showing greater aIns response to social betrayal in the Trust Game exhibited reduced aIns-vmPFC connectivity in response to social fairness in the Ultimatum Game.

**Figure 5.**
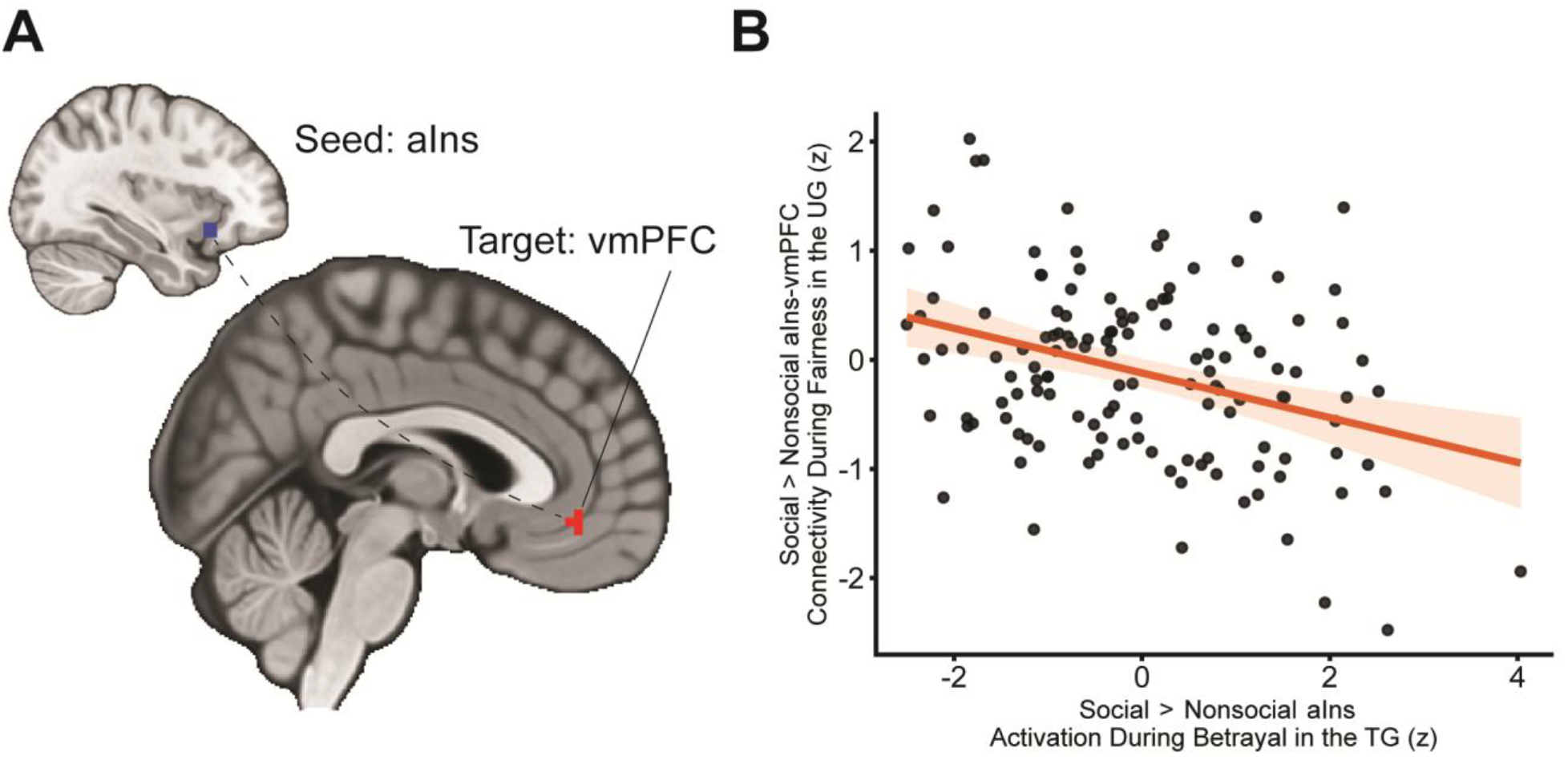
Whole-brain PPI analysis with anterior insula seed. **A.** A whole-brain psychophysiological interaction (PPI) analysis using the anterior insula (aIns) as the seed revealed a significant cluster in ventromedial prefrontal cortex (vmPFC). **B**. Scatterplot showing the negative association between Trust Game anterior insula activation and Ultimatum Game aIns-vmPFC connectivity.

## Discussion

Fairness and trust are often examined separately, but both depend on evaluating whether another agent has violated an expected social norm (Fehr & Gächter, 2002; Fehr & Camerer, 2007). The first result of the present study was behavioral: acceptance rates in the Ultimatum Game increased monotonically with offer fairness, replicating the core finding that responders are willing to sacrifice monetary gain to resist inequitable allocations (Güth et al., 1982; Bolton & Zwick, 1995; Henrich et al., 2001; Sanfey et al., 2003). More importantly for the present study, participants did not show a reliable social-versus-nonsocial difference in acceptance behavior. The behavioral results therefore establish that participants treated the UG as a robust fairness task: acceptance increased with offer equity, whereas human and computer proposers did not differ reliably. That null partner effect matters because any neural social-versus-nonsocial effects are unlikely to be reducible to overt differences in acceptance behavior.

Against that behavioral backdrop, the clearest neural result was a negative parametric relationship between offer fairness and left anterior insula activity in the UG. This pattern closely matches prior UG work showing stronger insular responses to unfair or norm-violating offers, as well as meta-analytic evidence identifying the anterior insula as a consistent node in fairness-related decision making (Sanfey et al., 2003; Civai, 2013; Gabay et al., 2014; Feng et al., 2015). Within that literature, anterior insula activity is commonly interpreted as reflecting the affective salience and aversiveness of perceived injustice (Chang & Sanfey, 2013; Corradi-Dell’Acqua et al., 2013). In the present data, this region scaled with offer unfairness across four levels, consistent with graded sensitivity to inequity rather than a categorical response to partner type.

The most novel finding came from the cross-task psychophysiological interaction analysis. Because regional anterior insula activation did not correlate directly across tasks, this analysis tested whether cross-task convergence might instead be expressed in how anterior insula couples with valuation systems during fairness evaluation. Individuals who showed larger anterior insula responses to social betrayal in the TG exhibited reduced anterior insula-vmPFC connectivity during the social-versus-nonsocial fairness contrast in the UG. A concrete, albeit tentative, interpretation is that stronger anterior insula recruitment to betrayal indexes a more reactive or vigilant salience profile, and that during fairness evaluation these individuals engage vmPFC-based integration less strongly rather than more strongly. This reading fits models in which anterior insula helps flag socially aversive norm violations, whereas vmPFC contributes to normative valuation and the integration of affectively relevant choice signals (Chang & Sanfey, 2013; Baumgartner et al., 2011; Xiang et al., 2013). On this account, the negative direction of the association is informative: heightened betrayal sensitivity may bias some individuals toward salience detection and vigilance, and away from fuller valuation-based integration, when they judge socially framed fairness outcomes. This network-level interpretation also fits broader accounts of the social brain, in which social connectedness and disconnection are supported by interactions among anterior insula, reward circuitry, and medial prefrontal systems rather than by isolated regional responses (Kim & Sul, 2023).

A few limitations qualify this conclusion. Although the Ultimatum Game and Trust Game both involve responses to social norm violations, they differ in structure, incentives, and psychological demands; accordingly, the present findings are better interpreted as evidence for a partial relationship between fairness- and betrayal-related processes than as evidence for a single shared mechanism. We also did not collect post-task subjective measures of how participants’ impressions of their partners changed over time, limiting our ability to determine whether neural responses tracked stable partner representations, shifting expectations, or affective reactions to specific outcomes. Future work should use more closely matched task designs, independent anterior insula ROIs, and direct measures of partner appraisal and expectation updating to clarify the scope and robustness of this cross-task relationship.

These findings help explain the dissociation between behavior and neural measures. Behaviorally, participants treated the Ultimatum Game as a fairness task: acceptance increased monotonically with offer equity, whereas human and computer proposers did not differ reliably. Neurally, however, social context still related to variability in anterior insula function and anterior insula-vmPFC coupling, consistent with prior work showing that partner identity and social framing can shape internal valuation even when overt choices remain tightly constrained by payoff structure (McCabe et al., 2001; Delgado et al., 2005; Stanley et al., 2012). Relatedly, work on social influence suggests that social signals can alter subjective utility through ACC, insula, and vmPFC mechanisms, providing one route by which social information can be incorporated into valuation (Chung et al., 2015). By measuring both tasks in the same individuals, we show that the strongest bridge between betrayal and fairness is not a shared behavioral phenotype or a simple one-to-one regional activation correspondence but a relationship between betrayal-related anterior insula reactivity and fairness-related anterior insula-vmPFC coupling.

## Methods

### Participants and Recruitment

We collected data from 225 adults (age range 19–80 years) recruited from the Temple University community and the surrounding Philadelphia area. This sample was drawn from a larger parent project designed to examine social decision making across the adult lifespan; the present study represents one of several investigations using data from this project. Sample size and exclusion criteria were pre-registered on OSF: https://osf.io/z6m35. Eligible participants were native English speakers with normal or corrected-to-normal vision and no history of neurological or psychiatric illness. All participants completed a health screening questionnaire and standard MRI safety screening prior to enrollment. Participants were told that one of the trials would be selected at random and added to their bonus payment. They were also informed that proposer partners would receive payment based on that random trial’s outcome, with any money won either going directly to the social partner or being returned to a pool of laboratory funds for outcomes with nonsocial partners. The study protocol was approved by the Temple University Institutional Review Board, and all participants provided written informed consent.

For the present analyses, we excluded participants over the age of 55 to minimize the risk of mild cognitive impairment and to focus on young and middle-aged adults. Participants were also excluded as outliers for demonstrating poor neuroimaging data quality (as assessed with MRIQC). We removed runs using a boxplot threshold applied to the tsnr and fd_mean IQMs from MRIQC. Specifically, outlier runs were defined as runs with fd_mean values exceeding Q3 + 1.5 × IQR, as well as those with tsnr values lower than Q1 − 1.5 × IQR, where Q1 and Q3 are the 25th and 75th percentiles, respectively, and IQR = Q3 − Q1. We note that the tsnr for the multi-echo data were derived from echo-2 to increase compatibility with the single-echo data. In addition, runs where participants missed more than 25% of task trials were excluded. After all the exclusions, data from 132 participants were included in the analysis. Table 1 provides a summary of the demographics for these participants.

**Table 1.**
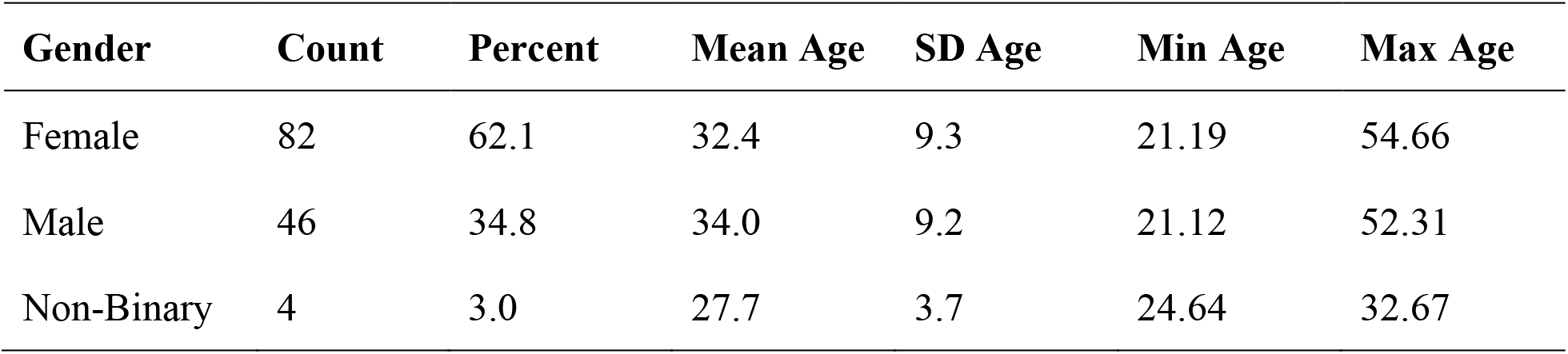
Participant Demographics by Gender.

| **Gender** | **Count** | **Percent** | **Mean Age** | **SD Age** | **Min Age** | **Max Age** |
| --- | --- | --- | --- | --- | --- | --- |
| Female | 82 | 62.1 | 32.4 | 9.3 | 21.19 | 54.66 |
| Male | 46 | 34.8 | 34.0 | 9.2 | 21.12 | 52.31 |
| Non-Binary | 4 | 3.0 | 27.7 | 3.7 | 24.64 | 32.67 |

### Experimental Procedure

Data collection occurred over two visits. During Visit 1 (≈ 90 min), participants underwent a mock MRI scan to acclimate to scanner noise and the supine position. Visit 2 (≈ 120 min), scheduled within one week of Visit 1, began with practice trials of both the Trust Game (TG) and Ultimatum Game (UG) on a laptop to minimize missing responses, followed by two fMRI runs, one TG and one UG, in counterbalanced order. After the scan, participants completed exploratory questionnaires.

### Tasks and Experimental Design Trust Game Task

On each trial, the participant (investor) received a maximum of $8 endowment (ranging from $0.00 to $8.00) and decided, in $1 increments, how much to send to a purported partner for that trial (Figure 1). The partner may either be their gender- and age-matched confederate (stranger), friend, or computer. Gender and age matching applied to the stranger condition only; friend partners were actual friends recruited by each participant, and computer partners involved no matching. For non-binary participants, the confederate’s gender was the participant’s assigned gender at birth. Sent amounts are tripled; the partner then either returns a portion (reciprocation) or returns $0 (betrayal). We modeled outcome-phase events as “reciprocated-trust” or “broken-trust,” each defined at the feedback onset. For analyses involving social > nonsocial comparisons, contrasts were specified as stranger > computer for better comparability to UG analyses, which only had computer and stranger conditions.

### Ultimatum Game (UG) Task

In the UG, participants are made offers by one of two different partners (a stranger or a computer) and decide whether to accept or reject the offer (Figure 1). When paired with a (social) stranger partner, participants were informed that their partner was a past participant that made real choices in past games and that their choices may similarly be used with future participants. Each stranger was experienced only once to maintain the one-shot design of the game. When paired with a (nonsocial) computer partner, participants were informed that their choices were decided by an algorithm, and the winnings would return to the general pool of lab funds. Participants are told that the partner received $16 or $32 and decided to share 5%, 10%, 25%, or 50% of this endowment with them. If they accept the offer, they both receive the money as the split proposes. However, if they reject the offer, neither the participant nor the partner will receive any money. Each trial consisted of the endowment phase (1 second), an interstimulus interval of (1 second), a decision phase of 3.25 s, and an intertrial interval of 1–7 s (M = 2.5 s).

### Neuroimaging Data Acquisition and Preprocessing

All images were collected using a 3T Siemens PRISMA MRI scanner (Siemens AG, Muenchen, Germany) with a 20-channel head coil at the Temple University Brain Research and Imaging Center (TUBRIC). Functional BOLD data were acquired with a multi-echo gradient echo-planar imaging (EPI) sequence with four echoes (13.8, 31.54, 49.28, 67.02 ms), simultaneous multislice acceleration (multiband factor = 3), in-plane acceleration (GRAPPA = 2), and partial Fourier set to 7/8. The sequence used a 80×80 matrix, 2.7 mm isotropic voxels, 10% slice gap, 51 axial slices, and TR = 1615 ms (Smith et al., 2024). Data were converted to BIDS using HeuDiConv (Halchenko et al., 2024), distortion-corrected using warpkIt (Van et al., 2023), and minimally preprocessed with fMRIPrep 24.1.1 (Esteban et al., 2019). Multi-echo denoising and TE-dependence analysis were performed using the TEDANA workflow (DuPre et al., 2021). Full preprocessing details appear in the Supplementary Methods.

### Neuroimaging Analysis

Below we outline the subject-level models and group-level contrasts used to test whether betrayal- and unfairness-related responses generalized across tasks. Analyses were conducted in FEAT (FSL v6.0.7; Jenkinson et al., 2012). For each participant, task regressors were convolved with a double-gamma hemodynamic response function. For the Trust Game, first-level models focused on the outcome phase (modeled as a fixed-duration epoch starting at the onset of the partner feedback window; 1 s) and included regressors for reciprocated and unreciprocated trust outcomes. For the Ultimatum Game, first-level models focused on the offer phase (modeled as a fixed-duration epoch starting at partner cue onset and ending at the fixed offset of the decision/response window; 5.25 s duration). Trials were modeled separately by partner type (social, nonsocial) and endowment level, with offer fairness included as a parametric modulator within each condition. This model allowed us to estimate partner-agnostic responses to unfairness as well as social-versus-nonsocial differences in fairness-related responses.

Our first-level UG activation model included 11 task-related regressors: four regressors modeled offer events separately for each endowment-by-partner condition (high-stranger, low-stranger, high-computer, low-computer); four regressors modeled the corresponding parametric modulation of offer events by offer fairness within those same conditions (high-stranger, low-stranger, high-computer, low-computer); two regressors modeled response-time effects (constant, parametrically modulated); and one regressor modeled missed trials. All task-related regressors were convolved with FSL’s canonical hemodynamic response function.

Our first-level TG activation model included 10 task-related regressors: three regressors modeled decision-phase events separately for each partner type (computer, friend, stranger); six regressors modeled outcome-phase events separately for each partner-by-outcome condition (computer-defect, computer-reciprocate, friend-defect, friend-reciprocate, stranger-defect, stranger-reciprocate); and one regressor modeled missed trials. All task-related regressors were convolved with FSL’s canonical hemodynamic response function.

First-level activation and connectivity models included nuisance regressors to account for motion and physiological noise. These included the six rigid-body motion parameters, the first six aCompCor components, two non-steady-state volumes, and framewise displacement. TEDANA-derived nuisance components were included as additional regressors to further account for residual non-BOLD variance. High-pass filtering was implemented with a 128 s cut-off using discrete cosine basis functions.

To identify regions sensitive to offer fairness, we conducted a whole-brain parametric modulation analysis of the UG offer phase, collapsing across partner type and endowment level. To identify regions sensitive to betrayal, we contrasted unreciprocated trust with reciprocated trust during the TG outcome phase. The anterior insula seed used in connectivity analyses was defined as the binary intersection of significant clusters from these two activation analyses: the UG fairness parametric modulation effect and the TG unreciprocated trust > reciprocated trust contrast.

To test task-modulated connectivity, we used a generalized psychophysiological interaction (gPPI) model centered on the anterior insula (McLaren et al., 2012). The physiological regressor was the time series extracted from the anterior insula seed. The model included the original UG task regressors, the anterior insula physiological regressor, and separate interaction terms formed by multiplying the mean-centered anterior insula time series by each task regressor. This gPPI model produced condition-specific estimates of anterior insula connectivity. Connectivity contrasts were then constructed to test whether anterior insula coupling varied as a function of the social relative to nonsocial fairness effect during the UG offer phase. The group-level model also included a between-subjects covariate representing each participants’ anterior insula response to social relative to nonsocial betrayal during the TG outcome phase. This tested whether individual differences in betrayal-related anterior insula activation predicted anterior insula connectivity during fairness evaluation in the UG.

Higher-level analyses used mixed-effects models implemented in FLAME (Woolrich et al., 2004). When multiple usable lower-level FEAT inputs were available for a participant, run-level estimates were combined using second-level fixed-effects models before group-level modeling. Group-level models also included covariates for age, gender, fd_mean, and tsnr. Whole-brain statistical maps were thresholded using a voxel-wise threshold of Z > 3.1 and cluster-extent correction at p < .05 (Worsley, 2001) based on Gaussian random field theory. Final statistical maps were visualized using MRIcroGL (Rorden, 2025).

### Deviations from Preregistration

During implementation of the preregistered analyses, we identified a minor discrepancy in the direction of the regression model for Hypothesis 1.1. The preregistered hypothesis stated that greater ROI activation to unreciprocated trust in the Trust Game would predict a higher tendency to reject unfair offers in the Ultimatum Game. However, the preregistered analysis plan reversed this direction, specifying a model in which rejection rates predicted ROI activation. To maintain conceptual consistency with the hypothesis, we followed the intended direction and modeled UG rejection tendencies as the outcome variable and TG ROI activation as the predictor. Additionally, our preregistration specified the dorsal anterior cingulate cortex (dACC) as the primary region of interest and the anterior insula as an additional region of interest for further investigation. In the analyses reported here, convergent effects across fairness processing in the UG and unreciprocated trust processing in the TG were observed in the anterior insula but not the dACC. The full set of preregistered results can be found in the Supplement. Accordingly, the present study focuses on the anterior insula. No other deviations from the preregistration were made.

## Acknowledgments

The authors gratefully acknowledge Cooper Sharp and Yi Yang for assistance with data analysis and feedback on preliminary results. This study was supported by National Institutes of Health grant RF1-AG067011.

## Data and Code Availability

Analysis code and selected derivative outputs are available on GitHub: https://github.com/DVS-Lab/rf1-betrayal.

